# An Engineered Genetic Circuit for Lactose Intolerance Alleviation Coupled with Gut Microbiota Recovery

**DOI:** 10.1101/2020.09.15.297564

**Authors:** Mingyue Cheng, Zhangyu Cheng, Yiyan Yu, Wangjie Liu, Ruihao Li, Zhenyi Guo, Jiyue Qin, Zhi Zeng, Lin Di, Yufeng Mo, Chunxiu Pan, Yuanhao Liang, Jinman Li, Yigang Tong, Yunjun Yan, Yi Zhan, Kang Ning

**Affiliations:** College of Life Science and Technology, Huazhong University of Science and Technology, 430074, Wuhan, P.R., China; Innovation Base of Life Science and Technology, Qiming College, Huazhong University of Science and Technology, 430074, Wuhan, P.R., China; Key Laboratory of Molecular Biophysics of the Ministry of Education, Huazhong University of Science and Technology, 430074, Wuhan, P.R., China; State Key Laboratory of Pathogen and Biosecurity, Beijing Institute of Microbiology and Epidemiology, 100071, Beijing, P.R., China; Beijing Advanced Innovation Center for Soft Matter Science and Engineering (BAIC-SM), College of Life Science and Technology, Beijing University of Chemical Technology, Beijing, 100029, Beijing, P.R., China

**Keywords:** Lactose intolerance, Genetic engineering, Synthetic biology, Gut microbiota, *In vitro* simulation, *In vivo* assessment

## Abstract

**Background:** Lactose malabsorption occurs in around 68% of the world populations, causing lactose intolerance (LI) symptoms such as abdominal pain, bloating and diarrhoea. To alleviate LI, previous studies mainly focused on strengthening intestinal β-galactosidase activity but neglected the inconspicuous colon pH drop caused by gut microbes’ fermentation on non-hydrolysed lactose. The colon pH drop will reduce intestinal β-galactosidase activity and influence the intestinal homeostasis.

**Results:** Here, we synthesized a tri-stable-switch circuit equipped with high β-galactosidase activity and pH rescue ability. This circuit can switch in functionality between expression of β-galactosidase and expression of l-lactate dehydrogenase in respond to intestinal lactose signal and intestinal pH signal. We confirmed the circuit functionality was efficient using 12-hrs *in vitro* culture at a range of pH levels, as well as 6-hrs *in vivo* simulations in mice colon. Moreover, another 21-days mice trial indicated that this circuit can recover lactose-effected gut microbiota of mice to the status (enterotypes) similar to that of mice without lactose intake.

**Conclusions:** Taken together, the tri-stable-switch circuit can serve as a promising prototype for LI symptoms relief, especially by flexibly adapting to environmental variation, stabilizing colon pH and restoring gut microbiota.

## Introduction

The lactose malabsorption, defined as the inefficient absorption of lactose was reported to has a global prevalence of 68%, which ranges from 28% in western, southern, and northern Europe to 64% in Asia and 70% in the Middle East [1]. The regional prevalence is extremely high in several countries requiring an efficient therapies such as 80% in Colombia (America), 85% in China (Asia), 98% in Armenia (Europe), 100% in South Korea (Asia), Yeman (Middle East), and Ghana (Sub-Saharan Africa) [1]. Lactose intolerance (LI) symptoms, defined as the presence of gastrointestinal symptoms caused by lactose malabsorption in the small intestine, will occur when non-hydrolysed lactose flows into the colon as the bacteria substrate [1, 2]. This non-hydrolysed lactose brings a high osmotic load into the colon luminal contents, which leads to increased water and electrolytes followed by softening stool, thus causing abdominal pain and cramps [3]. Meanwhile, this lactose can be fermented into lactic acid and other short chain fatty acids with gaseous products such as hydrogen, CH_4_, and carbon dioxide, thus causing flatulence and diarrhoea [3, 4].

The current treatments for LI mainly include dietary control, enzyme replacement and probiotic supplement. For dietary control, the moderation or restriction of lactose intake is recommended to relieve symptoms [5–7], which impacts people’s enjoyment of dairy products. Additionally, a recent study found that the administration of the highly purified short-chain galactooligosaccharide can help to adjust gut microbiome to improve the LI [8]. Enzyme replacement is another important approach for large populations of LI individuals [9]. The intake of exogenous lactase may help lactose digestion and absorption for LI subjects, but its efficacy still lacks convincing evidence [2]. Compared to short-acting enzyme replacement, probiotic supplements have an advantage in their sustainability [10] and a certain number of studies have confirmed that they can alleviate LI [11–13]. The key function of the probiotic is to enhance the intestinal β-galactosidase (β-GAL) activity of LI individuals for lactose digestion. In addition, the endogenous β-GAL produced by the probiotic is able to persist in a more stable manner in the intestine. However, the conventional bacteria cannot deal with the pH drop caused by fermentation of gut microbiota. The pH drop would not only cause physical discomfort such as diarrhoea, but also probably reduce the β-GAL activity [14, 15], and influence the intestinal homeostasis.

Genetical engineering, which can make the precise control over genome sequence [16], might be the solution for the pH drop problem faced by non-modified bacteria. Current designs of engineered bacteria have been confirmed as effective through the development of synthetic biology, for purposes such as infectious disease treatment [17] and cancer diagnostics [18]. Moreover, bacteria engineered by synthetic biology are believed to work more precisely and efficiently in addressing these diseases [19] compared to wild type bacteria. Previously, a recombinant starin expressing food-grade β-GAL for LI was constructed and evaluated [20, 21]. However, this engineered strain was still unable to deal with the pH drop. A stress-responsive system might make the bacteria more adaptable to the pH variation [22]. On the other hand, the influences of bacteria administration and pH drop on gut microbiota remain unclear. The influences might be understood by observing variations of gut microbiota during the lactose intake and bacteria administration phases.

In this study, we firstly designed and constructed a tri-stable-switch circuit using synthetic biology in the plasmid *pet-28a-1* with two functional states (accumulation of β-GAL and pH rescue) in response to signals of intestinal lactose and intestinal pH variation. Secondly, we transformed the circuit into the strain *Escherichia coli* BL21 (*E. coli* BL21) to form the engineered strain BL21: *pet-28a-1*, which was then used to confirm the circuit functionality *in vitro* and *in vivo*. Lastly, we investigated on variation of mice gut microbiota and found that the administration of engineered strain BL21: *pet-28a-1* can recover mice gut microbiota affected by excess lactose intake.

## Results

### The tri-stable-switch circuit can switch between two functionalities in response to environmental change

The tri-stable-switch circuit in the plasmid *pet-28a-1* (Fig. 1a, Additional file 2: Table S1 and Additional file 3) was designed based on a tri-stable switch derived from bacteriophage *lambda* [23]. The mutant lactose-inducible promoter placm (Additional file 2: Table S1) and pH-responsive promoter patp2 (Additional file 2: Table S1) were applied to sense the signal. The key enzymes applied within the system were the products of *lacZ* (β-galactosidase, β-GAL) and the fusion gene *ompA-lldD* (L-lactate dehydrogenase, L-LDH). To confirm the functionality of the circuit, we choose the strain *Escherichia coli* BL21 as the chassis which has been commonly used for stable expression of nontoxic exogenous protein. The circuit was transformed into the strain *Escherichia coli* BL21 (*E. coli* BL21) to form the engineered strain BL21: *pet-28a-1*. BL21: *pet-28a-1* was then able to dynamically switch between two functional states, which was regulated by lactose signal and pH signal: accumulation of β-GAL to digest lactose, and expression of L-LDH to rescue pH drop.

**Fig. 1.**
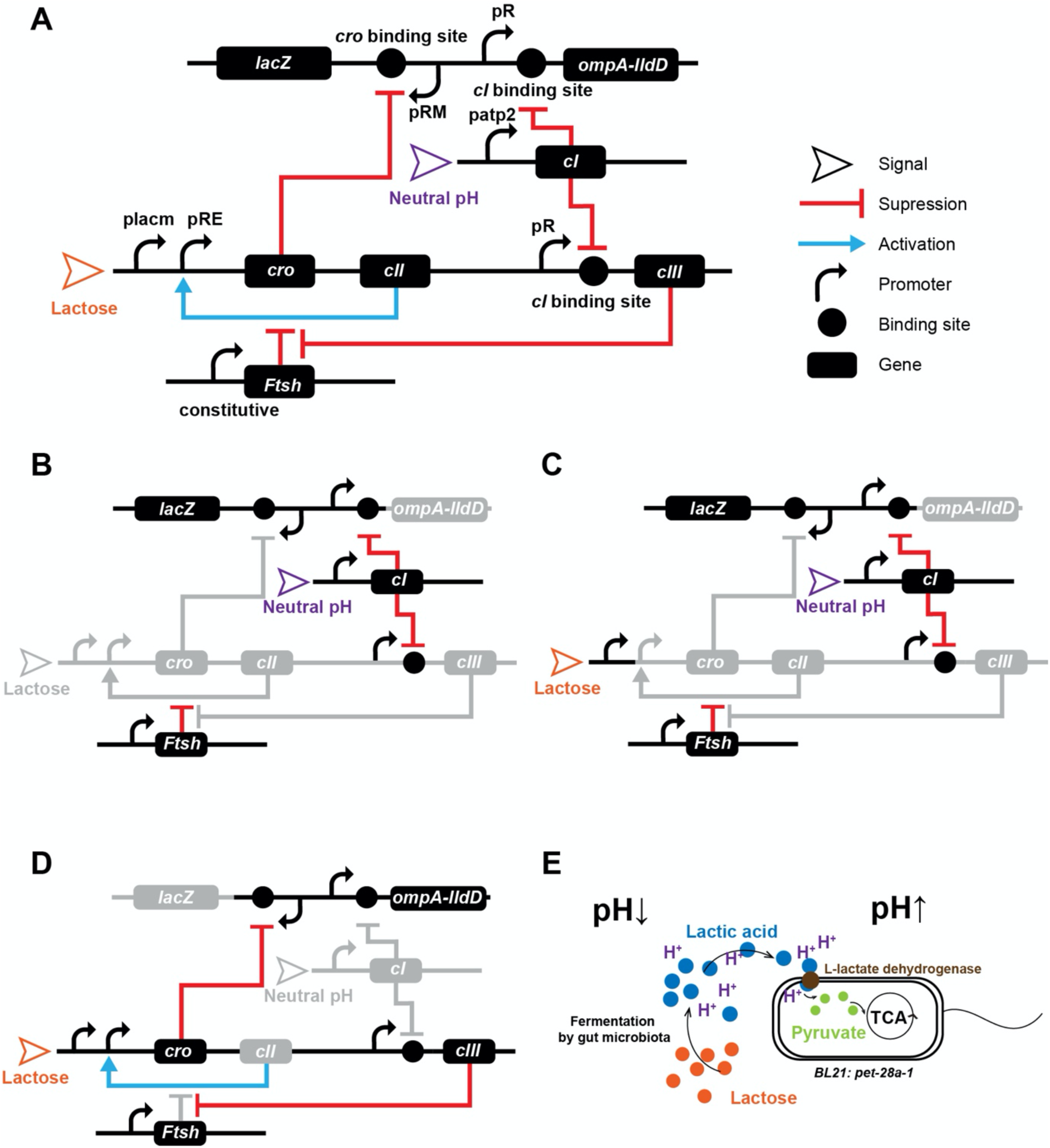
The tri-stable-switch circuit can switch between two functionalities in response to environmental change. **a** The design diagram of the tri-stable switch circuit. Parts of the circuit are derived from the bacteriophage *lambda*. The two promoters placm and patp2 are selected for sensing the lactose and pH signals, respectively. **b** When the BL21: *pet-28a-1* colonizes in the colon with a neutral pH, *lacZ* is stably expressed and accumulates β-GAL intracellularly. **c** When a flux of unabsorbed lactose occurs in the colon, the system switches to a transition state in response to lactose and pH signals. The expression of *ompA-lldD* for L-lactate dehydrogenase is strengthened and the expression of *lacZ* is weakened. **d** The system then focuses on expression of *ompA-lldD*. **e** The fermentation of lactose by gut microbiota causes pH drop, while the expressed L-lactate dehydrogenase can transform lactic acid into pyruvate, thus recovering the pH. The pyruvate then permeates into the cell for TCA pathway.

BL21: *pet-28a-1* can accumulate β-GAL after it colonized the colon (Fig. 1b). The mean pH of 7.0 in the human colon [24], as a signal, maintained continuous *cI* expression by inducing promoter patp2. The *cI* expression, which could hinder the transcripts of the gene after pR [25], then suppressed expression of *ompA-lldD* and *cIII*, thus ceasing the function of pH rescue. At this moment, the engineered bacteria would focus on the expression of the *lacZ* and accumulate β-GAL for supplementary lactose digestion when unabsorbed lactose fluxed into the colon.

BL21: *pet-28a-1* would gradually switch from *lacZ* expression to *ompA-lldD* expression after lactose fluxed into the colon (Fig. 1c). On the one hand, the lactose as a signal would trigger the promoter placm, thus activating the positive feedback loop of pRE, *cro* and *cII*. The *cro* expression then began to suppress the *lacZ* expression after the pRM through binding to its binding site [25]. Additionally, the *cro* expression can be strengthened by the *cII* expression. Nevertheless, owing to the suppression on the *cII* expression by endogenous *Ftsh* expression [26], this strengthening was suppressed to a certain degree. On the other hand, fermentation of lactose by gut microbiota would produce lactic acid and other short chain fatty acids, leading to acute drop of pH in the colon. This pH drop would weaken the patp2, suppressing the *cI* expression. However, the previous expressed products of *cI* would still suppress the expression of *ompA-lldD* and *cIII* to a certain degree, the suppression of which would gradually diminish because of degradation of these products. Hence the *ompA-lldD* expression would gradually recover to a normal condition, producing a signal peptide [27] andL-LDH [28, 29], which would be translocated on the cell membrane to transform lactic acid to pyruvate in the periplasm. In the meantime, the gradually recovered *cIII* expression would eliminate the suppression on *cII* expression by suppressing endogenous expressed *Ftsh* [26]. The unsuppressed *cII* expression then strengthened the *cro* expression, thus accelerating the cease of *lacZ* expression. Together, at the beginning of the period when lactose fluxed into the colon, the whole system was in an intermediate state of double functions. This was caused by the signals brought activation and deactivation of the pathways as well as the remained products such as repressor proteins which were not timely degraded.

Once these products were degraded, the BL21: *pet-28a-1* would then focus on *ompA-lldD* expression (Fig. 1d). The suppression on expression of *cIII* and *ompA-lldD* would be removed. On the one hand, constitutively expression of *cIII* eliminated the suppression on *cII* expression by endogenous expressed *Ftsh*, thus keeping lactose-activated positive feedback loop working to cease the *lacZ* expression. On the other hand, the *ompA-lldD* expression kept producing efficient signal peptides andL-LDH to transform lactic acid to pyruvate to rescue the pH drop (Fig. 1e). Afterwards, the pyruvate was transported into the cell by its carrier protein [30, 31] for the tricarboxylic acid (TCA) cycle [32]. Together, in this round of lactose intake, the engineered bacteria finished the lactose digestion as well as pH rescue.

BL21: *pet-28a-1* would subsequently switch to β-GAL accumulation for the next round of lactose ingestion. In this way, the BL21: *pet-28a-1* would switch back and forth in response to pulsed lactose intake.

### The tri-stable-switch circuit was efficient under a range of pH simulation *in vitro*

The promoters and transcriptional factors of the circuit have been tested by fluorescence detection (Additional file 1: Supplementary methods, Additional file 2: Table S2 and Table S3). We then tested whether the whole circuit can work effectively *in vitro* and *in vivo*. In the *in vitro* experiment (Fig. 2), we prepared mediums of three pH sets with 0.1% lactic acid and 1% lactose (Additional file 1: Supplementary methods) to simulate the pH range of acidic conditions caused by excess lactose intake in the human colon, whose normal pH is around 7 [24], as well as the mice colon, whose normal pH is around 5 [33]. Three pH sets included pH set I (initial pH = 4.54 ± 0.012), pH set II (initial pH = 5.34 ± 0.02), and pH set III (initial pH = 6.25 ± 0.02).

**Fig. 2.**
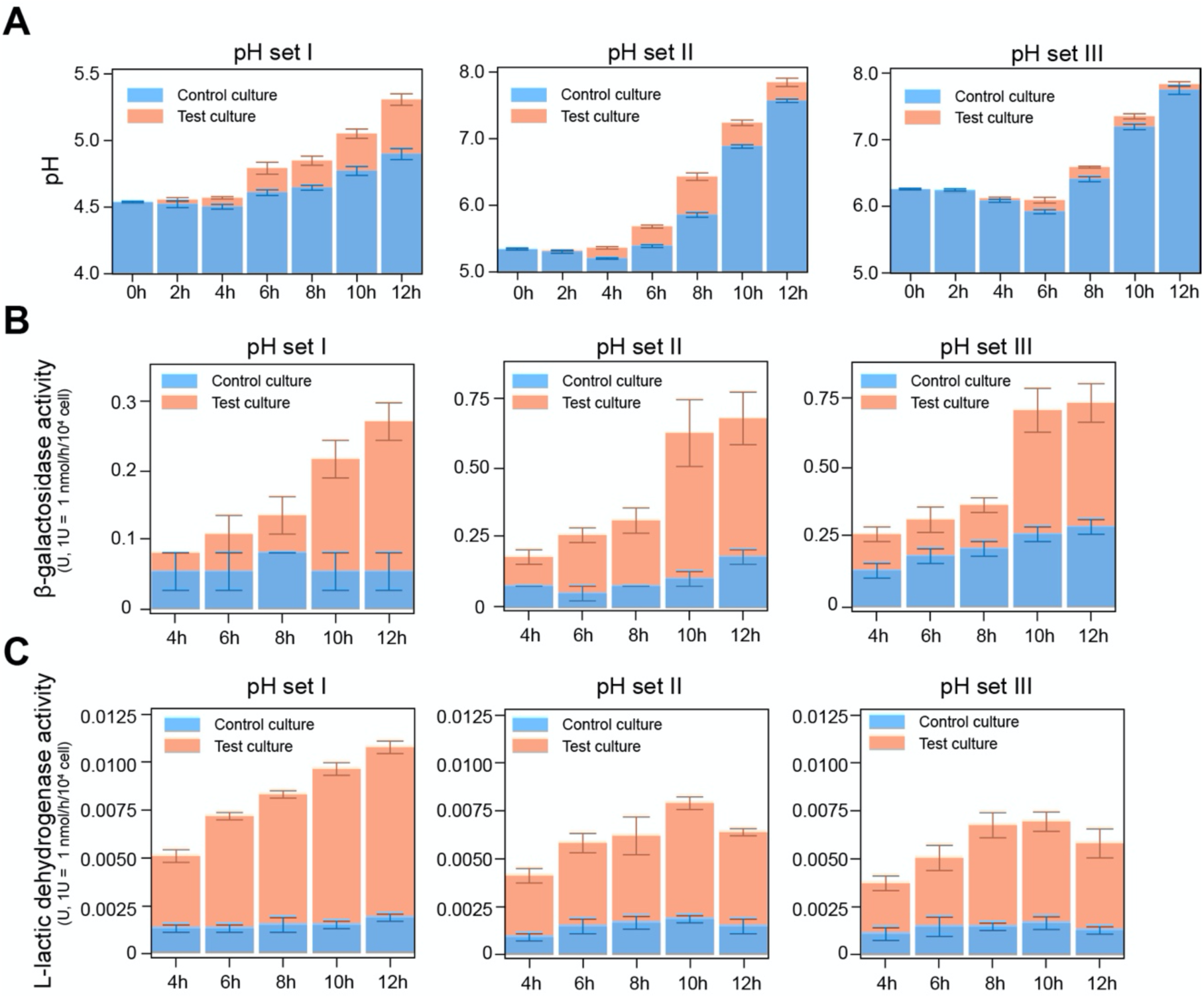
The tri-stable-switch circuit was efficient under a range of pH simulation *in vitro*. **a** The mean ± s.e.m. of pH variation during 12 hrs of the cultures of the bacteria grown at different set of pH. **b** The mean ± s.e.m. of β-GAL activity during 12 hrs of the cultures of the bacteria grown at different set of pH. **c** The mean ± s.e.m. of l-lactate dehydrogenase activity during 12 hrs of the cultures of the bacteria grown at different set of pH. In all panels, the control culture (BL21: *pet-28a-0*) is colored in orange, while the test culture (BL21: *pet-28a-1*) is colored in blue.

We subsequently recorded the variation of pH values and the expressed enzyme activity (Additional file 2: Table S5) during the following 12-hrs culturing of the bacteria at three pH levels, including the control strain (BL21: *pet-28a-0*) and the test strain (BL21: *pet-28a-1*). In addition, the control strain BL21: *pet-28a-0* (Additional file 4) was the *E. coli* BL21 transformed with empty vector. As was shown in Fig. 2a, pH values of the control culture and the test culture begin to increase at 6h. The increase of pH in the control culture was mainly associated with two processes including the metabolism of the massive increase of the bacteria and the consumption of the medium, while there was another additional process in the test culture that the expressed L-LDH helped to digest the lactic acid to increase pH. The additionally increased pH caused by L-LDH process was evident in the pH set I: The pH of the test culture increased to a higher degree than that of the control culture (test culture: 4.54 ± 0.02 to 5.31 ± 0.075; control culture: 4.54 ± 0.01 to 4.9 ± 0.072). The additionally increased pH in the test culture was observed in the pH set II and III as well, though not as obvious as that in the pH set I.

As was shown in Fig. 2b and c, the β-GAL activity and L-LDH activity of the test group were higher than those of the control group, which was caused by the expression of the *lacZ* gene and *ompA-lldD* gene of the circuit in the BL21: *pet-28a-1*. Before 4h, the measures for enzyme activity was unavailable because of the minimal amount of bacteria. After 4-hrs culturing, the β-GAL activity of the test group kept increasing stably in all three pH sets. In addition, from 8h to 10h in pH set II and pH set III, the β-GAL activity of the test group increased to the greatest extent and later flattened. In the other hand, the L-LDH activity of test group began to decrease in pH set II and pH set III after 10-hrs culturing. The corresponding pH range of the test group during 8h to 10h was 6.43 ± 0.10 to 7.23 ± 0.07 in pH set II, and 6.58 ± 0.03 to 7.34 ± 0.07 in pH set III, which meant that the switch of the functionality of the circuit was completed in this pH range. These results suggested that the relatively low pH promoted the L-LDH expression of the circuit to remove the lactic acid for increasing pH, and the increased pH can make the circuit begin to switch gradually from L-LDH expression to β-GAL expression, which would be completed in a pH range close to a neutral condition.

### The tri-stable-switch circuit helped mice to recover the pH drop caused by excess lactose intake

The *in vitro* experiment has confirmed the theoretical feasibility of the tri-stable-switch circuit to alleviate LI by switching between β-GAL expression and L-LDH expression, while whether it can work in *in vivo* still remained unclear. We thus recruited 84 mice, divided into five groups including the Initial set (n = 4), the Untreated group (n = 20), the Model group (n = 20), the Control group (n = 20), and the Test group (n = 20) to investigate on how the circuit worked *in vivo*. As was shown in Fig. 3a, in the first one week, mice in the Control group and the Test group were daily administrated with bacteria (BL21: *pet-28a-0* in the Control group, BL21: *pet-28a-1* in the Test group. OD_600_ = 1) in a total volume of 0.3 mL 0.9% NS suspension. The bacteria have been confirmed to colonize the mice colon, which can last at least 24 hrs (Additional file 1: Supplementary methods). Other groups were daily administrated with the same volume of NS. At 0h, mice of the Initial set were killed to test colon pH values as the pH value at 0h for all groups, and mice of other four groups were administrated with the lactose solution (12 mg of lactose per 20 g of body weight). In the following six hrs, the pH values of mice colons of the remained four groups were tested at each time point (Additional file 2: Table S6), and then plotted in Fig. 3b to show the pH variation. It was observed that the colon pH value of the Model group and the Control group evidently decreased to 4.66 ± 0.15 and 4.72 ± 0.25 from 0h to 3h, respectively, and then recovered to 4.89 ± 0.24 and 4.94 ± 0.1 from 0h to 3h, respectively. However, the colon pH value of the Untreated group without lactose intake, and the colon pH of the Test group using BL21: *pet-28a-1* kept relatively stable. Notably, during 2h to 4h, more colon lactose in Test mice was transformed to glucose rather than lactate, as compared to Model mice (Additional file 1: Supplementary methods). Therefore, these results indicate that the tri-stable-switch circuit is able to prevent the mice colon from pH drop caused by excess lactose intake, helping to keep intestinal homeostasis and relieve LI.

**Fig. 3.**
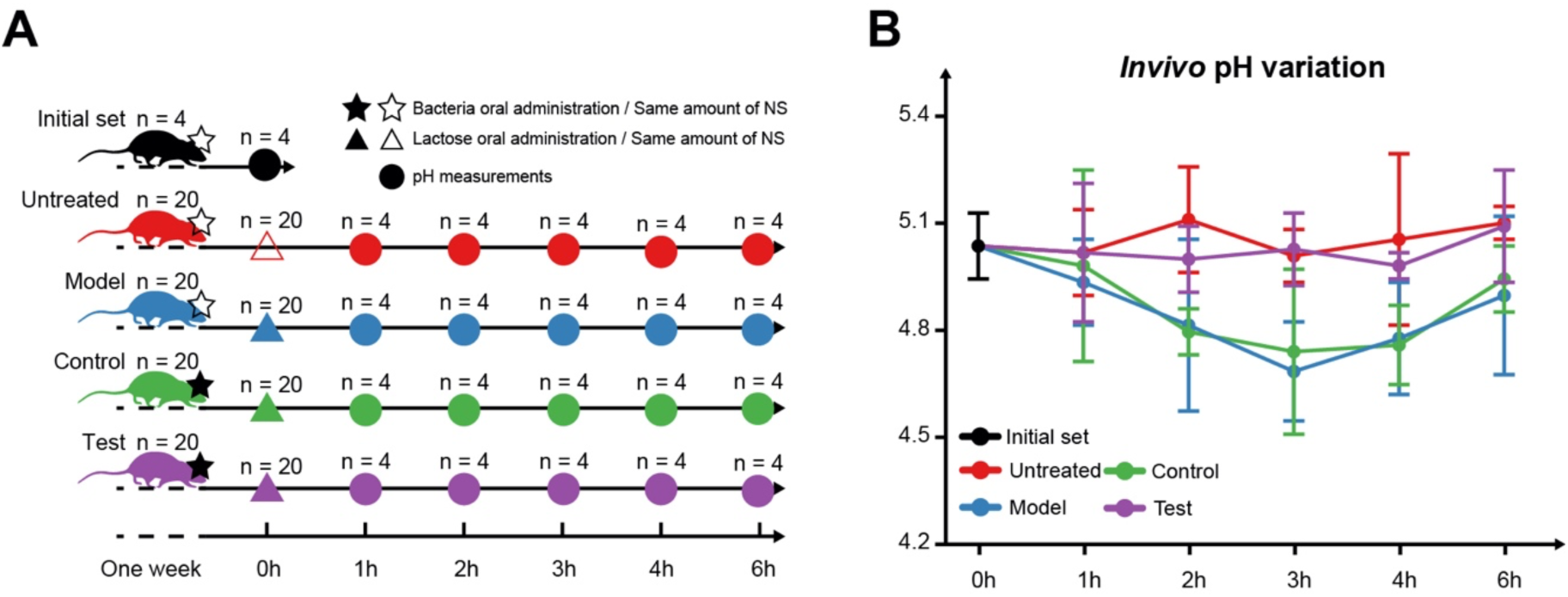
The tri-stable-switch circuit helped mice to recover the pH drop caused by excess lactose intake. **a** Five groups of mice including the Initial set (n = 4), the Untreated group (n = 20), the Control group (n = 20), the Test group (n = 20) were firstly subjected to different operations in one week. Mice in the Control group and the Test group were daily administrated with bacteria (BL21: *pet-28a-0* in the Control group, BL21: *pet-28a-1* in the Test group. OD_600_ = 1) in a total volume of 0.3 mL 0.9% NS suspension. Other groups were daily administrated with the same volume of 0.9% NS. At the time point of 0h, mice of the Initial set were killed for pH measures, and mice of other four groups were administrated with the lactose solution (12 mg of lactose per 20 g of body weight). In the following 6 hrs, four mice of each group were killed at each time point for pH measures. **b** The mean ± s.d. of pH variation of the mice colon during 6 hrs. The Initial set is designated as the initial point of four other groups. The pH variation of different groups is coloured differently.

### The tri-stable-switch circuit helped mice gut microbiota recovery from the effects of excess lactose intake

To trace the effects of the engineered bacteria equipped with the tri-stable-switch circuit on mice gut microbiota, we conducted a time-series trial with the high-frequency sampling of mice fecal samples (Additional file 2: Table S7). As was shown in Fig. 4a, four groups of mice (the Untreated group, the Model group, the Control group and the Test group) were under different interventions. The trial lasted for 21 days, divided into the four phases: normal care (Phase I), lactose challenge (Phase II), bacterial treatment (Phase III) and restoration (Phase IV). For Phase I, during which the four groups received normal care, the objective was to stabilize the physical signs and gut microbiota of mice in the four groups. For Phase II, during which lactose was fed to the Model, Control and Test mice, the objective was to investigate the influence of excess lactose. Phase III, in which the BL21: *pet-28a-1* was fed to the Test group while empty-vector-containing BL21: *pet-28a-0* used for mice in the Control group, was used to determine whether the BL21: *pet-28a-1* can alleviate LI. In Phase IV, we intended to observe whether the bacteria caused any side effects in the host mice.

**Fig. 4.**
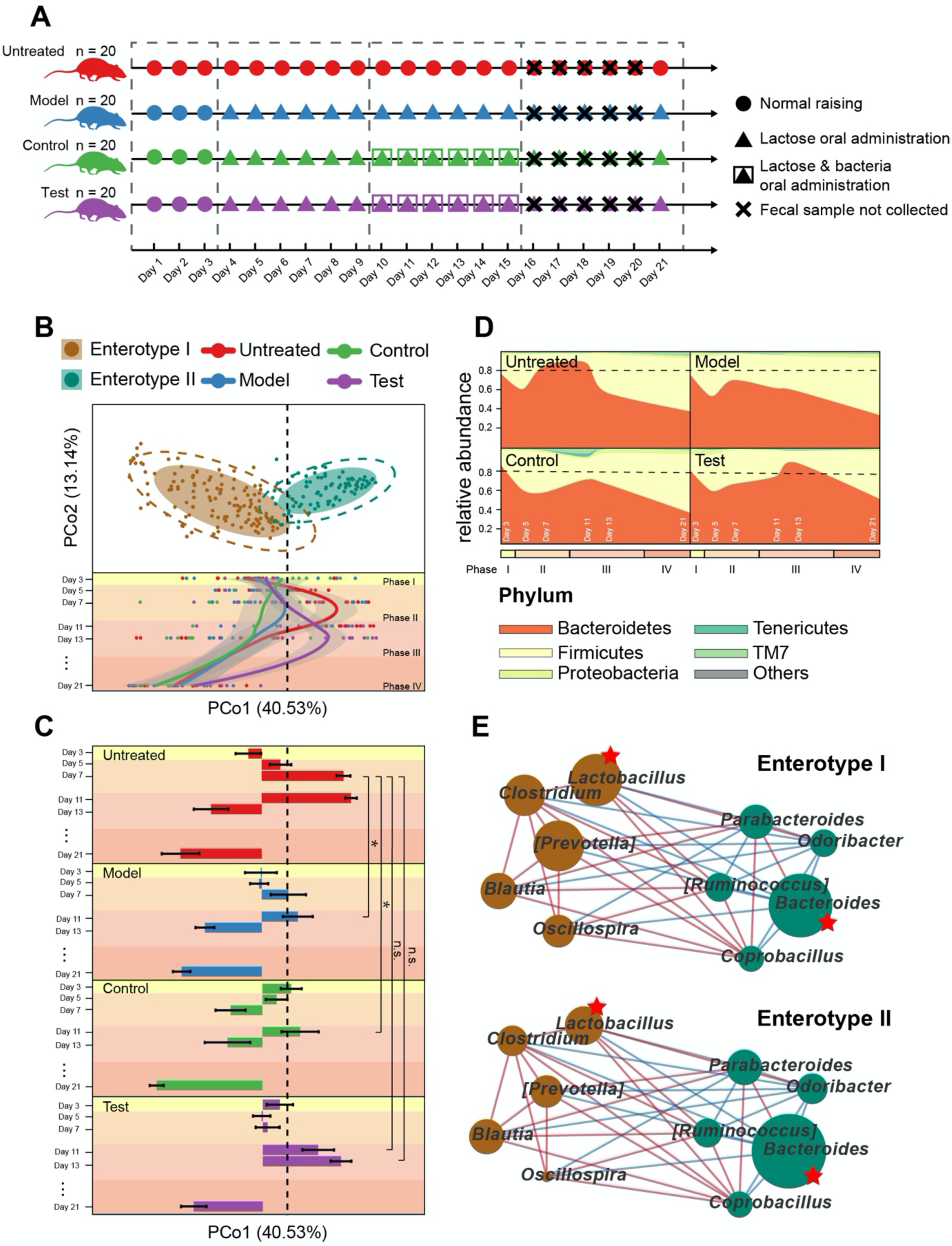
The tri-stable-switch circuit helped mice gut microbiota recovery from the effects of excess lactose intake. **a** The design of the mice trial for gut microbiota profiling (Detailed operations are available in the Additional file 1: Supplementary methods). **b** Top panel: Individual mice gut microbiota composition in the Untreated group (54 samples), Model group (55 samples), Control group (53 samples), and Test group (59 samples), stratified into two enterotypes and plotted on a JSD PCoA plot. The shaded ellipses represent an 80% confidence interval. The dotted ellipse borders represent a 95% confidence interval. Bottom panel: The gut microbiota samples are plotted according to their collection date on the y axis over 21 days, and their position on the x axis is plotted according to their first principal coordinate in the JSD-based PCoA (top panel). A Loess regression is applied to these points using the collection date and PCo1 coordinates, and the curves are plotted in different colors according to their groups, with a 95% pointwise confidence interval band in the gray shade. The dashed line is plotted to divide the area of the two enterotypes. **c** The mean ± s.e.m. of PCo1 coordinates from the four trial groups across 21 days. The dashed line represents the position on the x axis that divides the areas of the two enterotypes. A delayed shift to Enterotype II was observed in the Test group. *P < 0.05, **P < 0.01, ***P < 0.001; ns, not significant. Mann-Whitney-Wilcoxon Test. **d** The taxonomic variation of the mean relative abundance at the phylum level among four groups over the 21 days are shown in a stream graph. The dashed line represents a mean relative abundance of 0.8. **e** The network is constructed using the top ten abundant genera, based on the Kendall correlation with p value < 0.5 (Kendall’s *tau* statistic test) and q value < 0.5 (Benjamini and Hochberg corrections). The size of the node represents the mean abundance among enterotype I samples (top panel) or enterotype II samples (bottom panel). The color of the node represents the enterotype where the genus is more abundant. The asterisk refers to the most discriminant genus. The color of the edge represents the positive (red) or negative (blue) correlation.

Assigning enterotypes is a way to describe and differentiate the variance of gut microbiota, by stratifying gut samples according to their microbial composition [34]. Two enterotypes were firstly identified using all the samples of mice gut microbiota (Fig. 4b), which were statistically validated by CH and SI index (Additional file 2: Table S9). Two enterotypes were evidently different in microbial composition at the taxonomic levels of phylum, class, order, family, and genus (Additional file 2: Table S10). Compared to the samples of enterotype I, the samples of enterotype II displayed lower bacterial richness (*P* value < 2.20 × 10^−16^, Mann-Whitney-Wilcoxon test). Additionally, in the lower-richness enterotype II, the proportion of Firmicutes was reduced (*P* value = 1.60 × 10^−13^, Mann-Whitney-Wilcoxon test), while the proportion of Bacteroidetes was increased (*P* value = 1.16 × 10^15^, Mann-Whitney-Wilcoxon test). The characteristics of these identified mice enterotypes were consistent with those reported in a previous study [35].

More interestingly, the dynamics of the mice gut microbiota differed among the four trial groups over the 21-day (four phases) trial (Fig .4b and c). During Phase I, the microbiota of the gut samples from all the groups were of the enterotype I. At days 5 and 7 (Phase II: lactose challenge), most of the microbiota in the gut samples from the Untreated group trended towards the enterotype II, while the ones in the other groups under lactose treatment mostly remained in enterotype I. On days 11 and 13 (Phase III: bacteria treatment), the microbiota in samples from the Untreated group still trended towards enterotype II, while the ones in the Model and Control groups were restricted to enterotype I. However, the microbiota of the gut samples from the Test group using the BL21: *pet-28a-1* in this phase were mostly in the enterotype II area, with an obvious time lag observed between the Test and Untreated groups. All the microbiota in the gut samples from the four groups finally turned back to enterotype I after normal care during Phase IV. These different dynamic patterns fit well with the data from the mouse weight index recorded during the trials, that the bacteria-treated mice were rescued from LI-induced weight loss (Additional file 1: Supplementary methods, Additional file 2: Table S8). Taken together, these results indicated that the engineered bacteria was able to rescue the gut microbiota in the Test group to the patterns similar to those of the Untreated group.

Moreover, at the phylum level (Fig. 4d), when lactose was administrated to mice during Phase II, the proportion of Bacteroidetes was reduced in the Model, Control and Test groups. Nevertheless, during Phase III, the abundance of Bacteroidetes in the Test group was rescued to reach the same level (> 0.8) as the one in the Untreated group during Phase II. Furthermore, a variation pattern at genus level of the Test group was observed to be similar to that of the Untreated group after a time lag as well (Additional file 1: Supplementary methods). We then displayed the network of the top 10 abundant genera (Fig. 4e). The genus *Lactobacillus*, as well as the genus *Bacteroides* was the most abundant (Additional file 2: Table S10) and the most discriminant biomarker (Additional file 1: Supplementary methods) of the enterotype I and enterotype II, respectively. These two genera were observed to be negatively correlated (*P* value = 1.90 × 10^−10^, Kendall’s *tau* statistic). Therefore, the growth of the genus *Lactobacillus*, induced by the excess lactose intake might play a role in inhibiting the growth of the genus *Bacteroides* and the dynamic switch to enterotype II in the Phase II and III, which should have happened in a normal condition as observed in the Untreated group. The administration of the BL21: *pet-28a-1* helped the mice of the Test group to remove this inhibition so that their gut microbiota can continue to proceed the enterotype switch in Phase III, just like the mice gut microbiota of the Untreated group have done in Phase II.

## Discussion

In this study, we designed a tri-stable-switch circuit with ability of β-GAL accumulation and pH rescue. The engineered bacteria equipped with this circuit can flexibly adapt to the variation of intestinal environment, thus timely digesting lactose and rescuing the intestinal pH drop, along with the recovery of gut microbiota affected by excess lactose intake. We believe using engineered bacteria equipped with this tri-stable-switch circuit can serve as a promising therapeutic method for LI.

The tri-stable-switch circuit makes up the defect of non-modified bacteria by not only digesting lactose, but also enabling an additional function of pH rescue. The pH drop caused by fermentation of gut microbiota would not only cause diarrhoea, but also reduce the activity of the intestinal β-GAL and the intestinal homeostasis. Therefore, the tri-stable-switch circuit was designed to response to the signals of pH and lactose concentration, and then dynamically switch between two functional states: accumulation of β-GAL and pH rescue. The accumulation of β-GAL can help to digest lactose with high efficiency; meanwhile the pH rescue function helps to keep the intestinal homeostasis, giving engineered bacteria with this circuit better adaptability to intestinal environment than non-modified bacteria. These two functions have been confirmed in our *in vitro* and *in vivo* test.

Interestingly, in a 21-days *in vivo* mice trial, the engineered bacteria with the tri-stable-switch circuit was demonstrated to be able to recover the mice gut microbiota from the effects of excess lactose intake. During the trial, mice gut microbiota was influenced and could provide feedback to the environmental changes: The fermentation of unabsorbed lactose by gut microbiota produced acids and gas, leading to pH drop in the colon. This change of the environment in the colon would affect the gut microbiota in return. On the other hand, colonization of engineered bacteria affected the microbiota through microbial interactions, and its expressed products would also influence the intestinal environment. To explore the patterns of gut microbiota variation, we firstly identified two enterotypes based on the stratification of mice gut microbiota, and then we used the boundary of these two enterotypes as a threshold to identify the degree of variation in the microbiota. The most significant difference between the Untreated groups and other groups was found during Phase II (lactose challenge): when most of the mice microbiota samples in the Untreated group trended towards the enterotype II area, the ones from other groups seemed to be suppressed in the enterotype I area. This suppression might be caused by the intake of excess lactose. However, the suppression on the Test samples was subsequently removed by the feeding of our engineered bacteria during Phase III (bacteria treatment).

This study also has limitations. First, this study mainly underscores the design of the tri-stable-switch circuit and the confirmation of its functionality. Hence, for this purpose, the used *Escherichia coli* BL21 strain would be proper for functionality confirmation of a prototype rather than for the therapeutic intention. To apply this tri-stable-switch circuit to human still needs more sophisticated studies to find a proper chassis and ensure safety. Second, it has already been realized that the pH of the mice colon is lower than that of human colon. The switch of the tri-stable-switch circuit is completed in a pH range close to a normal condition, which better fits in human intestine. Nevertheless, in the *in vitro* experiment under a broad range of pH set, we have observed that it can still work well under a lower pH condition during the gradual process of the switch. Thirdly, the 16S rRNA gene sequencing cannot classify some potentially important gut microbes to the species level, the strain level, and the gene level. Nevertheless, the observed macroscopic trend of the gut microbiota variation has confirmed the recovery effects of the circuit, though the explanation of this trend still needs further investigations.

## Methods

### Experimental design

The gene sequences of the tri-stable-switch circuit were firstly synthesized by Integrated DNA Technologies and assembled by 3A assembly. The 3A-assembled intermediate parts were then assembled using In-Fusion. The circuit was transformed into the *E. coli BL21* for the following *in vitro* and *in vivo* experiments. *In vitro* experiments were designed to investigate on the variation of pH value, variation of β-galactosidase (β-GAL) activity, and variation of l-lactate dehydrogenase (L-LDH) activity of the bacterial culture under different pH levels. *In vivo* experiments were designed to investigate on the variation of pH value of the mice colon after intake of excess lactose. In addition, the sets of mice with or without administration of the engineered bacteria were used to observe the ability of pH rescue of the tri-stable-switch circuit. Another 21-days *in vivo* experiment was used to investigate on the gut microbiota variation of the mice after intake of excess lactose. The sets of mice with or without administration of the engineered bacteria were used to observe the effects of the tri-stable-switch circuit on gut microbiota.

The details of the circuit construction such as 3A assembly and fluorescence detection, the details of the *in vitro* experiments such as media preparation and measurements of enzyme activity, and the details of the *in vivo* experiments such as mice operations, 16S rRNA gene sequencing and microbiome analysis are available in the Supplementary Materials and Methods.

### Statistical Analysis

For categorical metadata and enterotype comparisons, samples were pooled into bins (Enterotype I/Enterotype II, Day 3/Day 5/Day 7…, etc.) and significant features were identified using Mann-Whitney-Wilcoxon Test with Benjamini and Hochberg correction of *P* values.

## Additional files

**Additional file 1: Supplementary methods**

The detailed designs and supplementary results of experiments including circuit construction, tri-stable switch confirmation, tri-stable circuit modelling, the *in vitro* experiment, the *in vivo* experiment, and the gut microbiota detection.

**Additional file 2: Supplementary tables**

The supplementary table 1 to 10 used for the manuscript and additional file 1.

**Additional file 3: The plasmid profile of *pet-28a-1***.

**Additional file 4: The plasmid profile of *pet-28a-0***.

## Ethics approval and consent to participate

Not applicable

## Consent for publication

Not applicable

## Availability of data and material

The datasets generated and analyzed during the current study are available in the short read archive (SRA) section of National Center for Biotechnology Information, under accession SRP152703.

## Competing interests

The authors have declared no competing interests.

## Funding

This project was supported by grants from the Ministry of Science and Technology of People’s Republic of China (Grant No. 2018YFC0910502), the National Natural Science Foundation of China (Grant No. NSFC-31871334 and NSFC-31671374), the Teaching Research Program from Hubei Province of China (Grant No. 2016071), and the National Undergraduate Training Programs for Innovation and Entrepreneurship (Grant No. 201710487069) from HUST and the Ministry of Education of China.

## Authors’ contributions

MY. C., ZY. C., YJ. Y., Y. Z. and K. N. designed the experiments. JM. L. and YG. T. conducted the DNA extraction and sequencing. MY. C., ZY. C., YY. Y., WJ. L., ZY. G., JY. Q., Z. Z., L. D., YF. M., and RH. L. conducted the plasmid constructions, fluorescence detection, and data analysis. MY. C., ZY. C., YY. Y, CX. P., and YH. L. conducted the *in vitro* and *in vivo* experiments. MY. C., ZY. C., Y. Z. and K. N. wrote and revised the manuscript.

## Acknowledgements

We thank all that have provided assistance for iGEM team HUST-China in iGEM 2016.

## References

1. Storhaug CL, Fosse SK, Fadnes LT. Country, regional, and global estimates for lactose malabsorption in adults: A systematic review and meta-analysis. Lancet Gastroenterol Hepatol. 2017;2:738–46.

2. Fassio F, Facioni MS, Guagnini F. Lactose maldigestion, malabsorption, and intolerance: A comprehensive review with a focus on current management and future perspectives. Nutrients. 2018;10.

3. de Vrese M, Stegelmann A, Richter B, Fenselau S, Laue C, Schrezenmeir J. Probiotics--compensation for lactase insufficiency. Am J Clin Nutr. 2001;73:421s–9s.

4. Heyman M. Effect of lactic acid bacteria on diarrheal diseases. J Am Coll Nutr. 2000;19:137s–46s.

5. Bohmer CJ, Tuynman HA. The effect of a lactose-restricted diet in patients with a positive lactose tolerance test, earlier diagnosed as irritable bowel syndrome: A 5-year follow-up study. Eur J Gastroenterol Hepatol. 2001;13:941–4.

6. Shaukat A, Levitt MD, Taylor BC, MacDonald R, Shamliyan TA, Kane RL, et al. Systematic review: Effective management strategies for lactose intolerance. Ann Intern Med. 2010;152:797–803.

7. Wilder-Smith CH, Olesen SS, Materna A, Drewes AM. Predictors of response to a low-fodmap diet in patients with functional gastrointestinal disorders and lactose or fructose intolerance. Aliment Pharmacol Ther. 2017;45:1094–106.

8. Azcarate-Peril MA, Ritter AJ, Savaiano D, Monteagudo-Mera A, Anderson C, Magness ST, et al. Impact of short-chain galactooligosaccharides on the gut microbiome of lactose-intolerant individuals. Proc Natl Acad Sci U S A. 2017;114:E367–e75.

9. Ianiro G, Pecere S, Giorgio V, Gasbarrini A, Cammarota G. Digestive enzyme supplementation in gastrointestinal diseases. Curr Drug Metab. 2016;17:187–93.

10. Ojetti V, Gigante G, Gabrielli M, Ainora ME, Mannocci A, Lauritano EC, et al. The effect of oral supplementation with lactobacillus reuteri or tilactase in lactose intolerant patients: Randomized trial. Eur Rev Med Pharmacol Sci. 2010;14:163–70.

11. Almeida CC, Lorena SL, Pavan CR, Akasaka HM, Mesquita MA. Beneficial effects of long-term consumption of a probiotic combination of lactobacillus casei shirota and bifidobacterium breve yakult may persist after suspension of therapy in lactose-intolerant patients. Nutr Clin Pract. 2012;27:247–51.

12. He T, Priebe MG, Zhong Y, Huang C, Harmsen HJ, Raangs GC, et al. Effects of yogurt and bifidobacteria supplementation on the colonic microbiota in lactose-intolerant subjects. J Appl Microbiol. 2008;104:595–604.

13. Oak SJ, Jha R. The effects of probiotics in lactose intolerance: A systematic review. Crit Rev Food Sci Nutr. 2019;59:1675–83.

14. Tomizawa M, Tsumaki K, Sone M. Characterization of the activity of beta-galactosidase from escherichia coli and drosophila melanogaster in fixed and non-fixed drosophila tissues. Biochim Open. 2016;3:1–7.

15. Juajun O, Nguyen TH, Maischberger T, Iqbal S, Haltrich D, Yamabhai M. Cloning, purification, and characterization of beta-galactosidase from bacillus licheniformis dsm 13. Appl Microbiol Biotechnol. 2011;89:645–54.

16. Hilton IB, Gersbach CA. Enabling functional genomics with genome engineering. Genome Res. 2015;25:1442–55.

17. Saeidi N, Wong CK, Lo TM, Nguyen HX, Ling H, Leong SS, et al. Engineering microbes to sense and eradicate pseudomonas aeruginosa, a human pathogen. Mol Syst Biol. 2011;7:521.

18. Danino T, Prindle A, Kwong GA, Skalak M, Li H, Allen K, et al. Programmable probiotics for detection of cancer in urine. Sci Transl Med. 2015;7:289ra84.

19. Hwang IY, Koh E, Wong A, March JC, Bentley WE, Lee YS, et al. Engineered probiotic escherichia coli can eliminate and prevent pseudomonas aeruginosa gut infection in animal models. Nat Commun. 2017;8:15028.

20. Li J, Zhang W, Wang C, Yu Q, Dai R, Pei X. Lactococcus lactis expressing food-grade beta-galactosidase alleviates lactose intolerance symptoms in post-weaning balb/c mice. Appl Microbiol Biotechnol. 2012;96:1499–506.

21. Zhang W, Wang C, Huang C, Yu Q, Liu H, Zhang C, et al. Construction and expression of food-grade beta-galactosidase gene in lactococcus lactis. Curr Microbiol. 2011;62:639–44.

22. Rajkumar AS, Liu G, Bergenholm D, Arsovska D, Kristensen M, Nielsen J, et al. Engineering of synthetic, stress-responsive yeast promoters. Nucleic Acids Res. 2016;44:e136.

23. Ptashne M. A genetic switch: Gene control and phage. Lambda. 1986.

24. Evans DF, Pye G, Bramley R, Clark AG, Dyson TJ, Hardcastle JD. Measurement of gastrointestinal ph profiles in normal ambulant human subjects. Gut. 1988;29:1035–41.

25. Schubert RA, Dodd IB, Egan JB, Shearwin KE. Cro’s role in the ci cro bistable switch is critical for {lambda}’s transition from lysogeny to lytic development. Genes Dev. 2007;21:2461–72.

26. Halder S, Datta AB, Parrack P. Probing the antiprotease activity of lambdaciii, an inhibitor of the escherichia coli metalloprotease hflb (ftsh). J Bacteriol. 2007;189:8130–8.

27. Pocanschi CL, Popot JL, Kleinschmidt JH. Folding and stability of outer membrane protein a (ompa) from escherichia coli in an amphipathic polymer, amphipol a8-35. Eur Biophys J. 2013;42:103–18.

28. Futai M, Kimura H. Inducible membrane-bound l-lactate dehydrogenase from escherichia coli. Purification and properties. J Biol Chem. 1977;252:5820–7.

29. Kimura H, Futai M. Effects of phospholipids on l-lactate dehydrogenase from membranes of escherichia coli. Activation and stabilization of the enzyme with phospholipids. J Biol Chem. 1978;253:1095–110.

30. Herzig S, Raemy E, Montessuit S, Veuthey JL, Zamboni N, Westermann B, et al. Identification and functional expression of the mitochondrial pyruvate carrier. Science. 2012;337:93–6.

31. Kristoficova I, Vilhena C, Behr S, Jung K. Btst, a novel and specific pyruvate/h(+) symporter in escherichia coli. J Bacteriol. 2018;200.

32. Deng Y, Ma N, Zhu K, Mao Y, Wei X, Zhao Y. Balancing the carbon flux distributions between the tca cycle and glyoxylate shunt to produce glycolate at high yield and titer in escherichia coli. Metab Eng. 2018;46:28–34.

33. McConnell EL, Basit AW, Murdan S. Measurements of rat and mouse gastrointestinal ph, fluid and lymphoid tissue, and implications for in-vivo experiments. J Pharm Pharmacol. 2008;60:63–70.

34. Arumugam M, Raes J, Pelletier E, Le Paslier D, Yamada T, Mende DR, et al. Enterotypes of the human gut microbiome. Nature. 2011;473:174–80.

35. Hildebrand F, Nguyen TL, Brinkman B, Yunta RG, Cauwe B, Vandenabeele P, et al. Inflammation-associated enterotypes, host genotype, cage and inter-individual effects drive gut microbiota variation in common laboratory mice. Genome Biol. 2013;14:R4.

